# DrugComb update: a more comprehensive drug sensitivity data repository and analysis portal

**DOI:** 10.1101/2021.03.25.436916

**Authors:** Shuyu Zheng, Jehad Aldahdooh, Tolou Shadbahr, Yinyin Wang, Dalal Aldahdooh, Jie Bao, Wenyu Wang, Jing Tang

## Abstract

Combinatorial therapies that target multiple pathways have shown great promises for treating complex diseases. DrugComb (https://drugcomb.org/) is a web-based portal for the deposition and analysis of drug combination screening datasets. Since its first release, DrugComb has received continuous updates on the coverage of data resources, as well as on the functionality of the web server to improve the analysis, visualization and interpretation of drug combination screens. Here we report significant updates of DrugComb, including: 1) manual curation and harmonization of more comprehensive drug combination and monotherapy screening data, not only for cancers but also for other diseases such as malaria and COVID-19; 2) enhanced algorithms for assessing the sensitivity and synergy of drug combinations; 3) network modelling tools to visualize the mechanisms of action of drugs or drug combinations for a given cancer sample; and 4) state-of-the-art machine learning models to predict drug combination sensitivity and synergy. These improvements have been provided with more user-friendly graphical interface and faster database infrastructure, which make DrugComb the most comprehensive web-based resources for the study of drug sensitivities for multiple diseases.

## INTRODUCTION

Despite the scientific advances in the understanding of complex diseases such as cancer, there remains a major gap between the vast knowledge of molecular biology and effective treatments. Next generation sequencing has revealed intrinsic heterogeneity across cancer samples, which partly explain why patients respond differently to the same therapy (1). For the patients that lack common oncogenic drivers, multi-targeted drug combinations are urgently needed, which shall block the emergence of drug resistance and therefore achieve sustainable efficacy (2). To facilitate the discovery of drug combination therapies, high-throughput drug screening techniques have been developed to allow for a large scale of drug combinations to be tested for their sensitivity and synergy in-vitro (3). Furthermore, patient-derived cancer cell cultures and xenograft models have been developed, which make the drug discovery closer to the actual patients (4–6).

With the increasing amount of drug sensitivity screening data, the challenge of translating them into actual drug discovery remains, as recent studies showed that most of clinically approved drug combinations work independently (7), that the efficacy and synergy observed in a pre-clinical setting may not be translated into a clinical trial (8). The challenge of utilizing the results from drug combination screens largely resides from un-harmonized metrics for synergy and sensitivity that are derived from different mathematical models, which are often incompatible for the same datasets (9). Another limitation is the lack of standardization of drug combination experimental design and the insufficient level of data curation and deposition to publicly available databases (10). Furthermore, the drug combination data has not been harmonized with single drug screening data, partially due to a lack of computational tools to enable a systematic comparison of drug combination efficacy against single drug efficacy (11).

To initialize the efforts for curating drug combination datasets, and to facilitate a community-driven standardization of evaluation of the degree of synergy and sensitivity of drug combinations, we have provided DrugComb as the very first data portal to harbour the manually curated datasets as well as the web server to analyse them (12). The original version of DrugComb consists of four major high-throughput studies, which served as a reference dataset for developing machine learning algorithms to predict drug combination sensitivity and synergy (13). Different from other recent databases including DrugCombdb (14) and SynergxDB (15), DrugComb is a unique resource as it is a compendium of database and web application, not only for depositing deeply curated public datasets but also for the analysis and annotation of user-uploaded data. Furthermore, DrugComb provides detailed visualization of drug combination sensitivity and synergy, which shall greatly facilitate the understanding of drug interactions at specific dose levels.

With the development of high-throughput screening techniques, the number of data points for drug combinations has been greatly increased. For example, the recent Dream Challenge on drug combination prediction has provided more than 20k drug combinations in cancer cell lines (16). Furthermore, drug combination screening has been extended to other disease models such as malaria and ebola (17). More recently, drug combination screening studies on COVID19 have been conducted, providing important clues for the treatment of the ongoing pandemic (18). In the new version of DrugComb, we aim to expand our manual curation from cancer to other diseases to improve the data coverage. On the other hand, drug combinations need to be harmonized with the monotherapy drug screening data, since these treatment options shall be evaluated using the same endpoint metric (such as progression free survival and overall survival) in clinical trials. Therefore, we aim to harmonize the drug combination with monotherapy drug screening, by providing informatics tools to evaluate their overall sensitivity in a more systematic manner. For this reason, in the new version of DrugComb, we do not limit ourselves for curating drug combination data, but rather we included monotherapy drug sensitivity screening data as well. More importantly, we provide a robust metric to enable a direct comparison of drug combinations and single drugs, as monotherapy drug screening can be considered as a subset of drug combination experiments. The new data harmonization framework thus allows a more systematic evaluation of a drug combination in comparison to a single drug. In addition, we implement several new modules for the analysis of these datasets, including the integration of drug targets and gene expressions of neighbouring proteins in a signalling network, such that the mechanisms of action of a drug or a drug combination can be annotated systematically in a specific cellular context. We also provide a baseline model based on CatBoost to predict the sensitivity and synergy of drug combinations, with which the machine learning community may develop novel algorithms to improve our understanding of drug responses in cancer cells. Taken together, the new version of DrugComb features an enhanced web portal to make drug screening data more interpretable and reusable for various applications such as machine learning, network modelling and experimental validation.

## RESULTS

### Overview of the DrugComb portal

DrugComb portal consists of two major components including a database for harbouring the most recent drug screening datasets as well as a web server to analyse and visualize these datasets or user-uploaded datasets for the degree of sensitivity and synergy. For retrieving the database, users can query by drug names, cell line names as well as study names. For utilizing the web server to analyse user-uploaded datasets, users need to import the data according to the format of an example file, and the results will be shown as both tabular and image displays, which are also downloadable. When users plan a drug combination experiment, they may utilize the web server to predict the sensitivity and synergy, and utilize such information to guide the selection of drugs. The drug targets as well as the gene expressions of the signalling pathways for a given cancer cell line can be also annotated as a network model. In the following, we describe how we have improved the coverage of the database as well as the data analysis modules of the web-server with a range of algorithms, and the new implementation techniques to accelerate data curation and harmonization efficiency (**Figure 1**).

**Figure 1.**
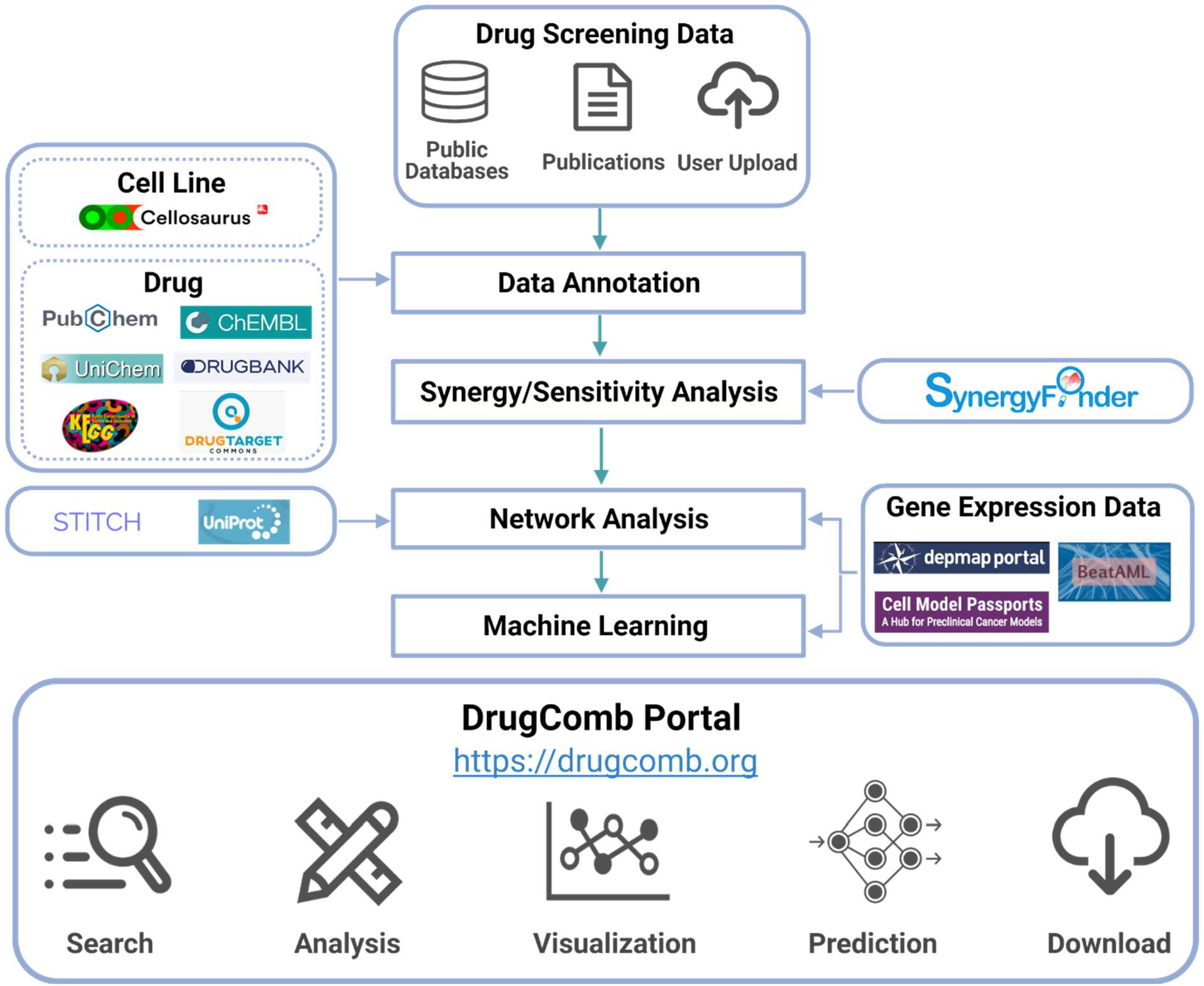
A schematic overview of the DrugComb database and web server pipeline. Drug combination and monotherapy drug screening datasets are curated from public databases, publications or user-upload. After quality control and pre-processing, the cell information is retrieved from Cellosaurus (19), while the drug information is retrieved from multiple databases including PubChem (20), ChEMBL (21), UniChem (22), DrugBank (23), KEGG (24) and DrugTargetCommons (25). The degree of synergy in drug combinations, as well as the sensitivity of drug combinations and single drugs are determined using the SynergyFinder R package (3). For inferring the mechanisms of action of drugs or drug combinations, their targets as well as interacting proteins are visualized in a signalling network, retrieved from STITCH (26) and UniProt (27). Furthermore, the gene expressions of these proteins in the given cancer cells are obtained from DepMap (28) and Cell Model Passports (29), and from BeatAML where the cancer samples were derived from AML (Acute Myeloid Leukaemia) patients (5). Machine learning algorithms utilize chemical structural and gene expression features to predict drug combination synergy and sensitivity. The DrugComb portal enables the query and download of curated raw datasets and analysis results, as well as the contribution of new datasets.

### Data sources

The initial version of DrugComb consists of four drug combination screening studies, covering 437, 923 drug combination experiments. We have curated much more drug combination experiments for cancer cell lines. Furthermore, we have incorporated monotherapy drug screening datasets and considered them as a subset of a drug combination experiment, where the other drug is absent. We have also included the drug screening results from patient-derived cancer samples in haematological malignancies (5). In addition to multiple cancer types, we have extended the curation efforts to other diseases such as ebola, malaria and COVID-19. The manual curation is under high level of quality control, that only those studies that reported the raw dose-response results will be considered, and thus the studies that reported only summary-level results including IC50, AUC (Area under the dose response curves) or synergy scores (e.g. combination index) are excluded. We have also standardized the metadata about experimental protocols of these studies so that their differences can be evaluated more systematically. The annotation of the bioassay protocols is based on the BAO (Bioassay annotation ontology) (30), that is commonly adopted for major chemical biology databases including ChEMBL (21), PubChem (20) and DrugTargetCommons (25). For the drugs and cell lines we provided the cross-database references such that their pharmacological and clinical information can be easily accessed (**Figure 2A-B**). As of March 2021, 751,498 drug combinations, 717,684 single drug screenings from 37 studies are deposited in DrugComb, corresponding to 21,621,279 unique data points spanning 2040 cell lines including 216 cancer types and three infectious diseases. **Table 1** shows the summary of the data points from the individual studies that are curated and harmonized in DrugComb.

**Figure 2.**
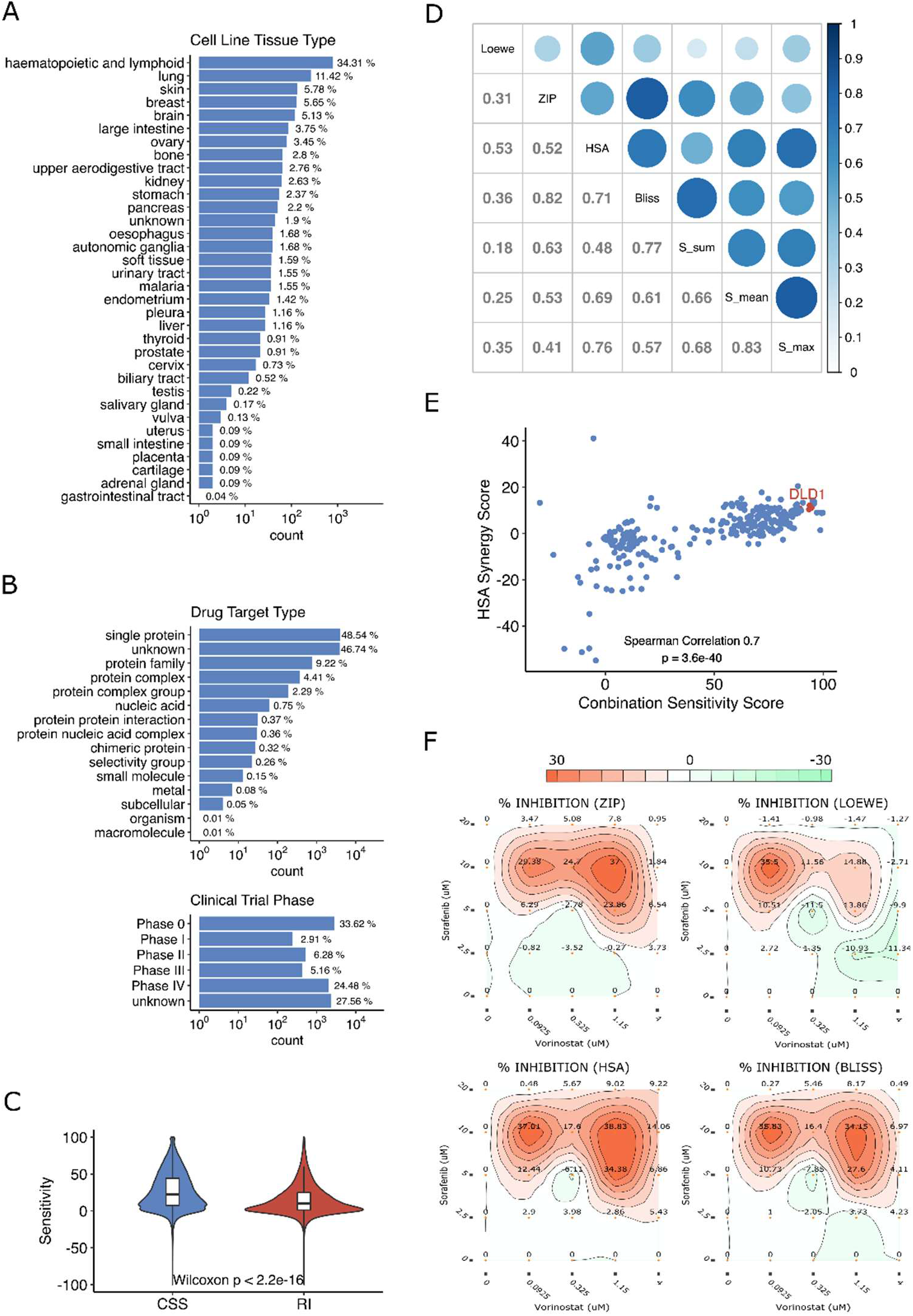
Overview of DrugComb data statistics. (A-B) Classifications of cell lines (n = 2040) and drugs (n = 8397). (C) The CSS score for drug combinations is higher than the RI score for monotherapy drugs, suggesting the general rationale for drug combination studies. (D) The correlations of synergy scores. (E) An example of SS plot for vorinostat and sorafenib combination across 128 cell lines. DLD-1 is a colon cancer cell line, which has shown strong synergy and sensitivity to the combination (34). (F) The synergy landscape over the dose-response matrix of vorinostat and sorafenib in DLD-1.

**Table 1.**
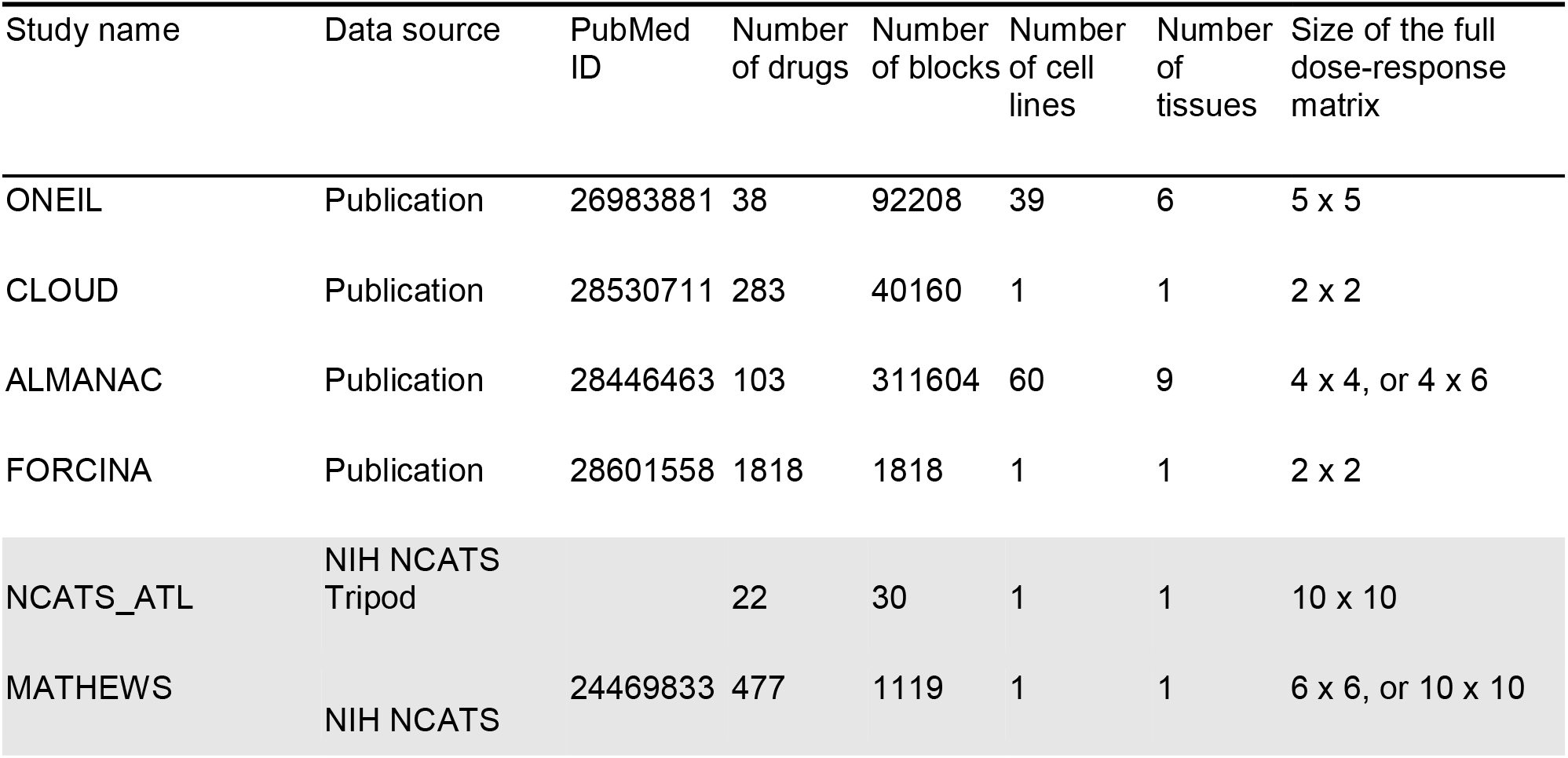
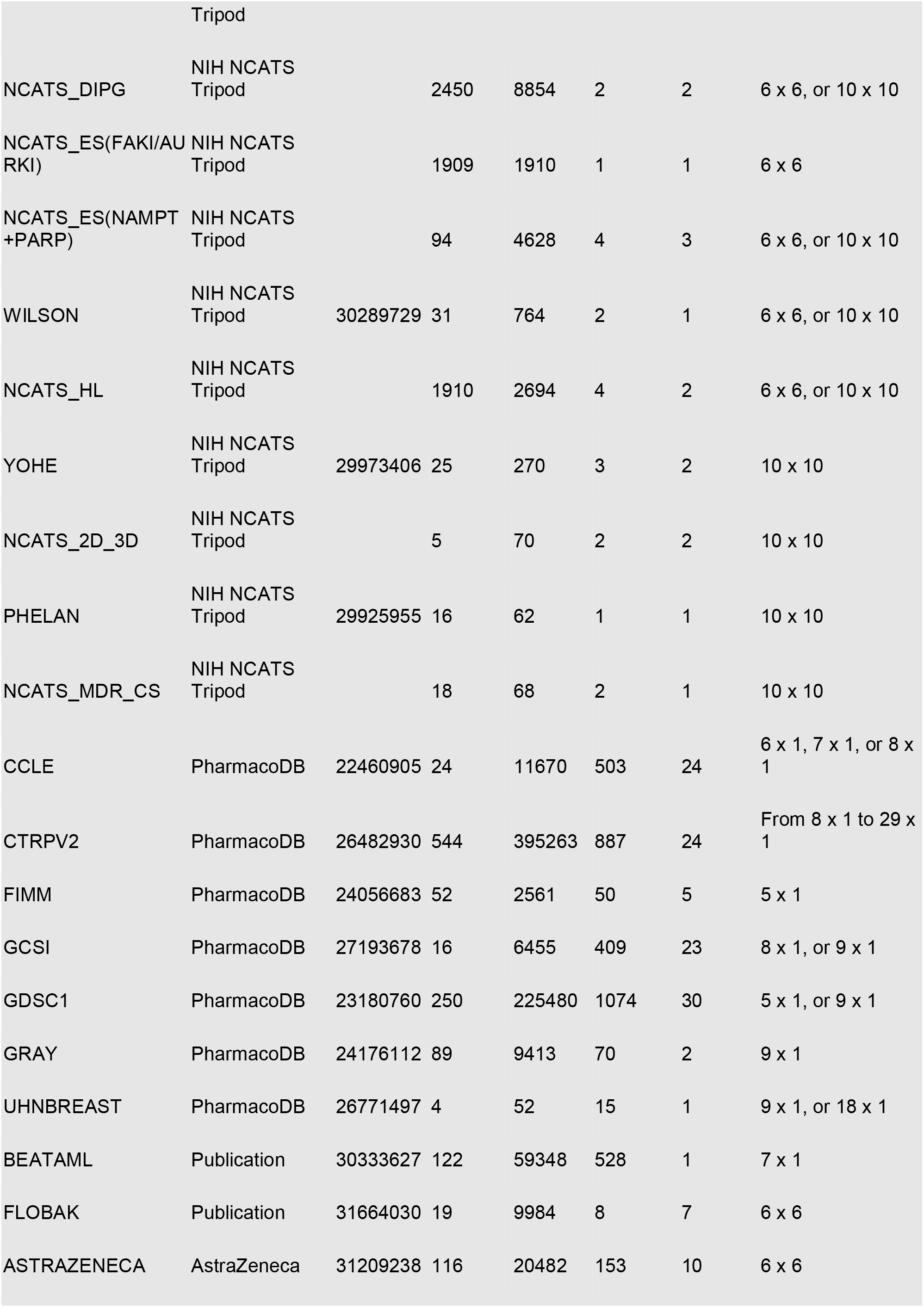
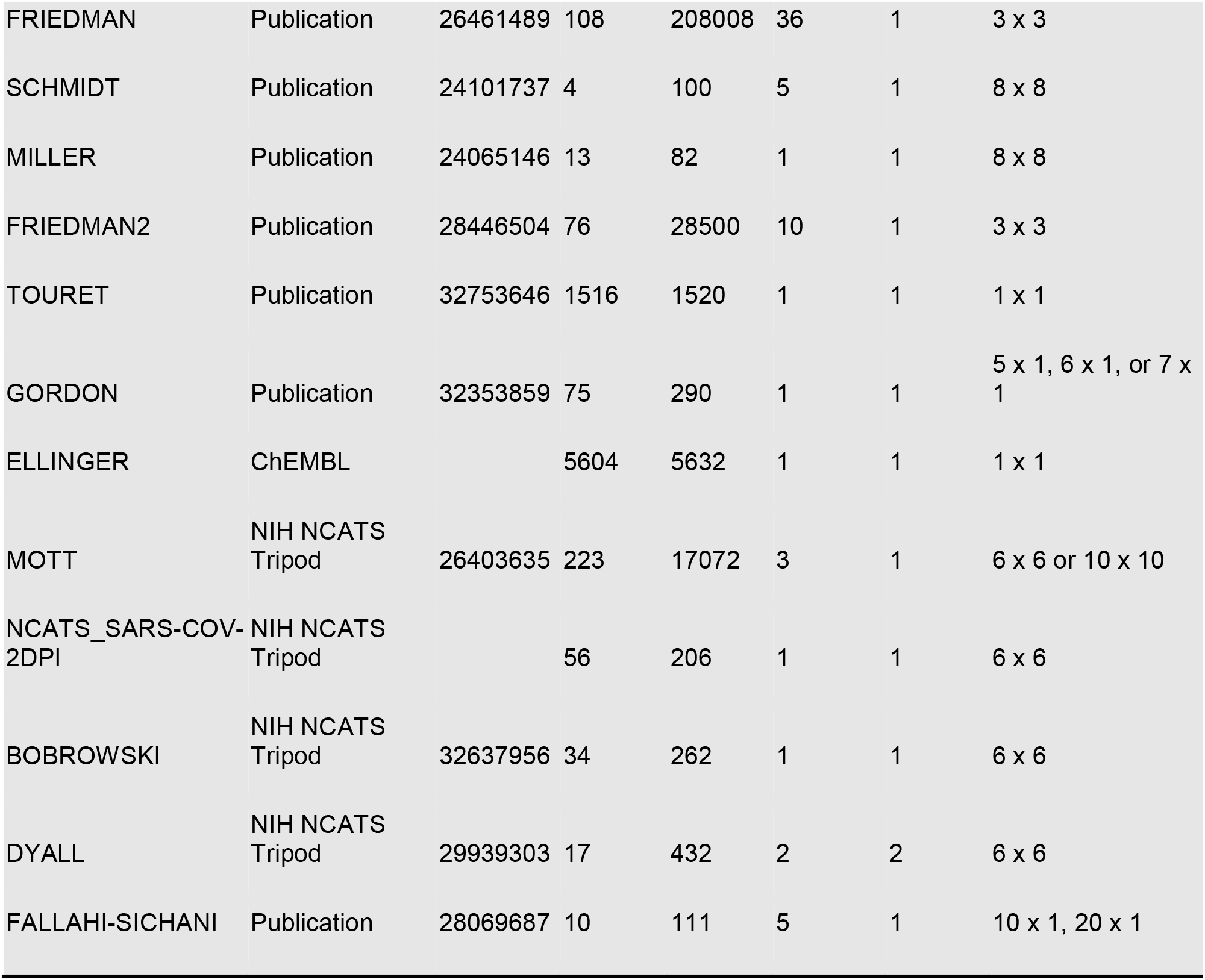
Database sources of DrugComb. Study names are determined by first authors of the publications or by the names of databases where the datasets were first deposited. Newly curated datasets are shown in the shaded areas as compared to the first four datasets that were deposited in the original version.

### Algorithms for assessing sensitivity and synergy

DrugComb utilizes the SynergyFinder R package to analyse drug combination sensitivity and synergy. The single drug sensitivity is characterized as a dose-response curve with its IC50 and RI (relative inhibition) values. RI is the normalized area under the log10-transformed dose-response curves, which has shown enhanced robustness to characterize drug sensitivity (31). Moreover, RI can be interpreted as percentage inhibition, summarizing the overall drug inhibition effects relative to positive controls. With the RI metric, drug responses of different concentration ranges can be compared, in contrast to IC50 or EC50, which are usually a relative term depending on the tested concentration ranges.

For drug combination sensitivity, we provide a metric called CSS (Combination Sensitivity Score), that is based on the normalized area under the log10-transformed of the combination dose-response curve when one of the two drugs is fixed at its IC50 concentration (11). CSS and RI use the same principle to characterize the overall drug response efficacy, such that their values can be directly compared (**Figure 2C**). For evaluating the drug synergy, we implement four major mathematical models including Bliss, Loewe, HSA and ZIP (32) and provide the visualization of these scores in the dose-response matrices. Furthermore, we provide a synergy score called S score that is derived from the difference between CSS and RI scores of the combination and single drugs respectively (11). The five synergy scores are based on different mathematical assumptions such that they do not necessarily match with each other (**Figure 2D**). For example, the Bliss model assumes probabilistic independence when drugs are non-interactive while the Loewe model assumes that the efficacy of non-synergistic drug combinations is identical to that of a drug combined with itself. The ZIP model, on the other hand, can be considered as an Ensembl model as it combines the assumptions of Bliss and Loewe (32). In actual clinical trials, approval of a drug combination often is based on the HSA model that simply shows that the drug combination improves patient survival compared to monotherapies. To insure the clinical translation of drug combinations, we encourage the use of all the major synergy scoring metrics, such that the top hits that pass the threshold of all of them can be prioritized (33). On the other hand, there have been biases by focusing solely on the synergy, while the sensitivity of a drug combination might be understudied. It is likely that a drug combination produces strong synergy while their overall efficacy is not achieving therapeutic relevance. Therefore, we provide an SS (Synergy-Sensitivity) plot to ensure that both of these two scores can be evenly weighted when interpreting the relevance of a drug combination (**Figure 2E, Supplementary Figure 1**).

As a unique feature of DrugComb, we visualize the synergy scores of a drug combination at each tested doses. The so-called synergy landscape allows a rich information display to facilitate the interpretation of the data, for which the most synergistic and antagonistic doses can be identified separately (**Figure 2F**). For a given drug or a given cell line, we provide the boxplots and histograms to show the general distributions of the synergy and sensitivity scores, such that the users may assess the general trend. For example, users may evaluate whether drug combinations involving a particular drug tend to be more synergistic, or a cell line tends to be more sensitive to drug treatment. Note that the majority of the data points (93.2%) that we curated from the literature do not contain replicates, and therefore, we decide not to provide the statistical significance of the synergy and sensitivity over a dose-response matrix, as the significance of individual doses contributing to the overall synergy cannot be systematically assessed. Therefore, we would like to highlight the issue of lack of replicates from a typical drug combination screening that may likely hinder the translation of the results into clinical trials.

### Network modelling for the mechanisms of action

Once a drug combination experiment has been conducted, for which the results were analysed with the sensitivity and synergy scoring, the next question would be the mechanisms of action of the drug combinations. Network modelling of drug combinations have been recently introduced as an efficient approach for the interpretation of drug combinations, as well as the identification of predictive biomarkers from molecular profiles of cancer (35–38). In DrugComb, the drugs are annotated with their target profiles, and these profiles were further annotated in the signalling networks of cancer cells, such that their first and secondary neighbour proteins can be also retrieved. We utilize the databases including ChEMBL, PubCHEM and DrugTargetCommons for their primary and secondary targets, and retrieve STITCH for the signalling networks. Furthermore, we have incorporated the transcriptomics profiles of the cancer cell lines into the network, such that their gene expression values can be also displayed (**Figure 3A**). In addition, we provide the correlation of the gene expression and drug sensitivity such that those neighbouring genes for which their gene expressions are highly correlated with the drug sensitivity will be further identified as potential biomarkers (**Figure 3B**).

**Figure 3.**
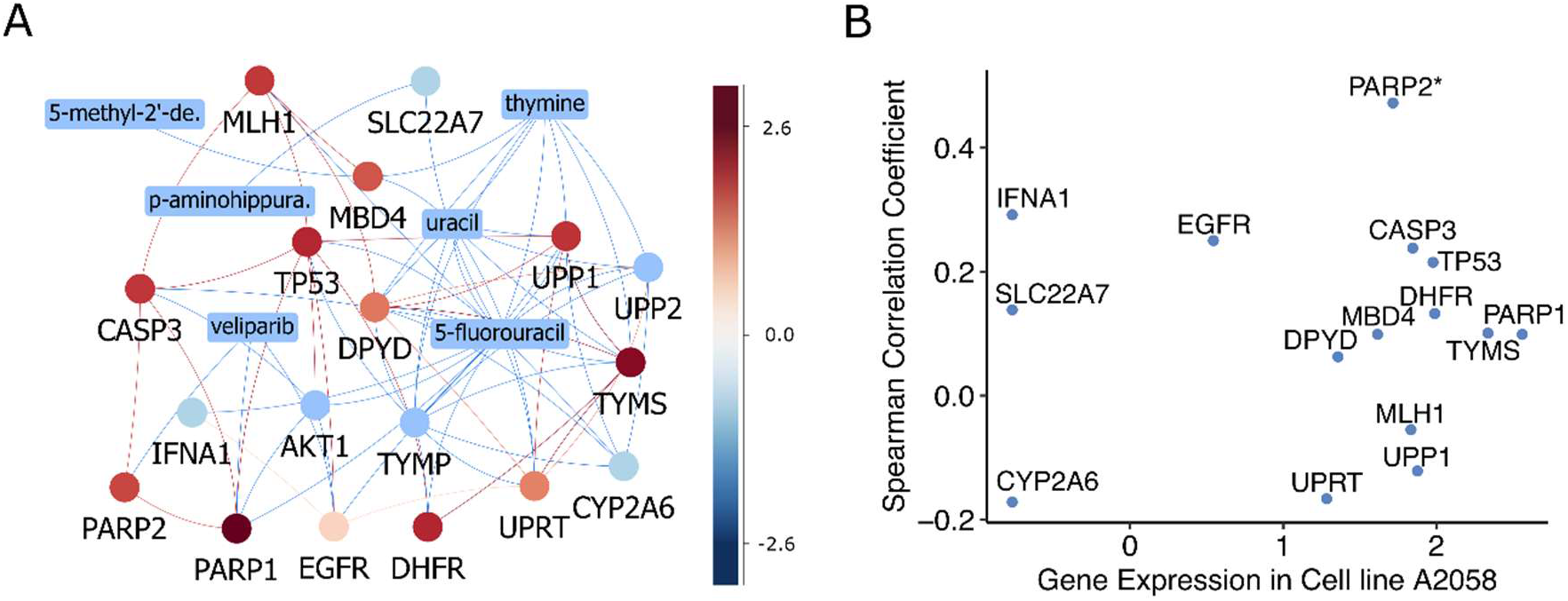
Network modelling of drug combinations. (A) An example of veliparib and 5-fluorouracil combination in A2058 melanoma cell line. Drug targets and neighbouring proteins are annotated with their gene expression values, some of which can be modulated by other drugs shown as rectangular boxes. (B) Gene expressions of the neighbouring proteins in A2058 as compared with the correlations of these genes with the drug combination sensitivity across all the tested cell lines. PARP2 is the primary target of veliparib, which shows top gene expression in A2058 as well as the highest correlation with the drug combination, suggesting that PARP2 is a potential biomarker for predicting the drug combination sensitivity of veliparib and 5-flurouracil.

For user-uploaded drug combinations or single drugs, ideally the InChiKeys of the drugs should be provided. This allows the web server to query drug STITCH ID from the major drug databases. In case only the drug names are provided, the web server will query from the major drug databases, for which their targets profiles will be visualized in a generic cancer signalling network. In case the cell line names can be matched with the existing gene expression data, their gene expression values will be displayed as coloured nodes. The network modelling results should be interpreted together with the actual drug screening profiles, such that the drug resistance or sensitivity can be related to its target or neighbouring gene expressions (**Figure 3B**).

### Machine learning for predicting sensitivity and synergy

Upon the large volume of drug combination data curated in DrugComb, we provide the state-of-the-art machine learning algorithms to predict the sensitivity and synergy for a user-selected drug combination on a given cancer cell line. We utilize the ONEIL data (39) to train a CatBoost model, which has been considered as a reference algorithm for many machine learning tasks (40). The ONEIL data consists of 583 drug combinations involving 38 drugs tested in 39 cell lines, resulting in 92,208 drug combination experiments consisting of 2,305,200 data points. The ONEIL data has been considered a high-quality dataset, as it contains multiple replicates and has been utilized in previous machine learning development (41–43). The CatBoost model is based on decision-trees that can facilitate the integration of different types of features including textual, categorical and numerical values. To build our model, the names of drugs and cell lines are specified as categories in our feature vectors. Additionally, the concentrations for drugs are considered as both numeric values and categories. The cell line’s gene expression and compound’s structural fingerprints (MACCS) are considered as numerical values. Moreover, in order to accelerate the training process for our model we consider only top 5% most variant genes (n = 153) across the 39 cell lines (**Figure 4**).

**Figure 4.**
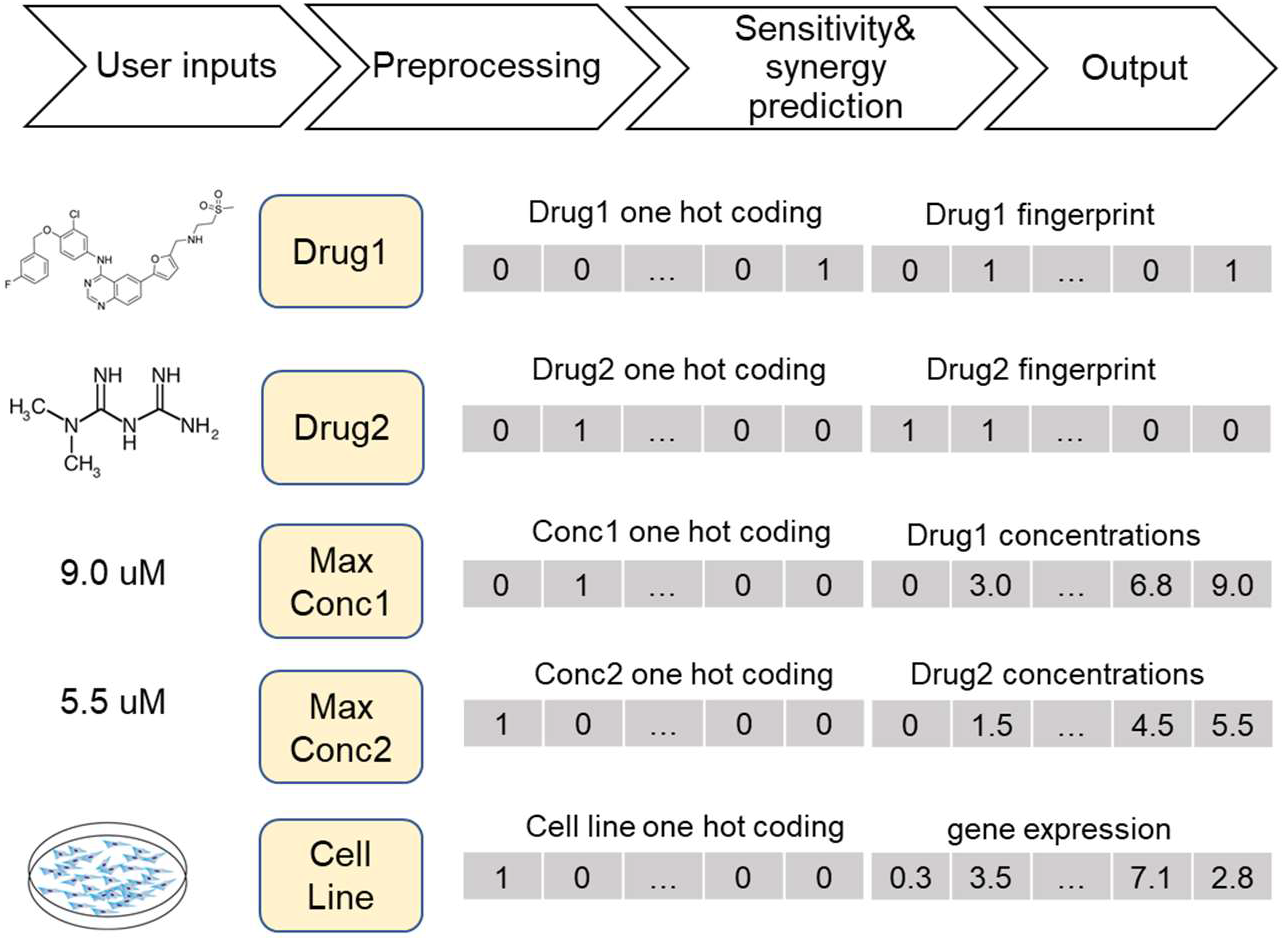
Work flow of machine learning prediction of drug combination sensitivity and synergy. Multiple features are integrated in CatBoost including one-hot encoding of drugs, concentrations and cell lines as well as their specific features including drug chemical fingerprints, drug concentrations and cell line gene expressions.

Among all the CatBoost hyper-parameters, only four of them show high importance for obtaining the best model. Those hyper-parameters include iterations that indicate the number of trees used in the model, maximum depth of the tree, the learning rate used for gradient steps, and the L2 regularization for the loss function. The best values for mentioned parameters are set and the rest of the hyper-parameters are set to the default values. For drug combination inhibition and synergy scores, a model has been trained separately and the results of the validation accuracy are presented in **Table 2**.

**Table 2.** Prediction accuracy of the CatBoost algorithm tested on ONEIL data.

|  | Correlation | R<sup>2</sup> | RMSE |
| --- | --- | --- | --- |
| INHIBITION | 0.98 | 0.97 | 7.12 |
| HSA | 0.79 | 0.62 | 8.03 |
| ZIP | 0.89 | 0.80 | 6.55 |
| LOEWE | 0.57 | 0.32 | 9.68 |
| BLISS | 0.87 | 0.75 | 7.65 |

To facilitate the prediction, users need only to specify the names and the maximal concentrations for each of two drugs, and a cell line name. After receiving the user input, the MACCS fingerprints of the drugs will be obtained by the RCDKlibs package in R, and the cell line gene expression data will be retrieved internally from DrugComb. The pre-processed data will be loaded into the trained models to predict the inhibition values and synergy scores for a 10×10 equally distanced dose matrix within the given maximal concentrations.

### Data contribution

To facilitate the data curation, we have provided a web server for users to upload their drug combination data into the database. The ‘Contribute’ panel will ask for the annotation information of the drug combination screening results, and then the actual data points will be formulated as a tabular format. We have utilized the contribution module to curate the majority of the literature datasets and found that it greatly facilitates the burden of the data contributors as well as data curators. For example, autofill functions are available when users input the literature citation and drug names. The cell line annotation is also available by retrieving the Cellosaurus website for its disease classification and other cross-reference links. Furthermore, data contributors are guided to provide critical information about assay protocols, such as detection technologies and culture time. When the data has been successfully uploaded, there are administrators who can view and approve the acceptance of the data, which will be shown as a separate study named by the first author of the publication (**Figure 5A**). In addition to the actual data points as an outcome of such a data curation effort, we will be able to systematically evaluate the differences in the assay protocols (**Figure 5B**), which might provide more insights on assessing the reproducibility of the drug sensitivity screens (44). Taken together, we believe that the data contribution may greatly facilitate the open access of drug screening data and therefore we encourage the users of DrugComb to be part of the community-driven data curation team.

**Figure 5.**
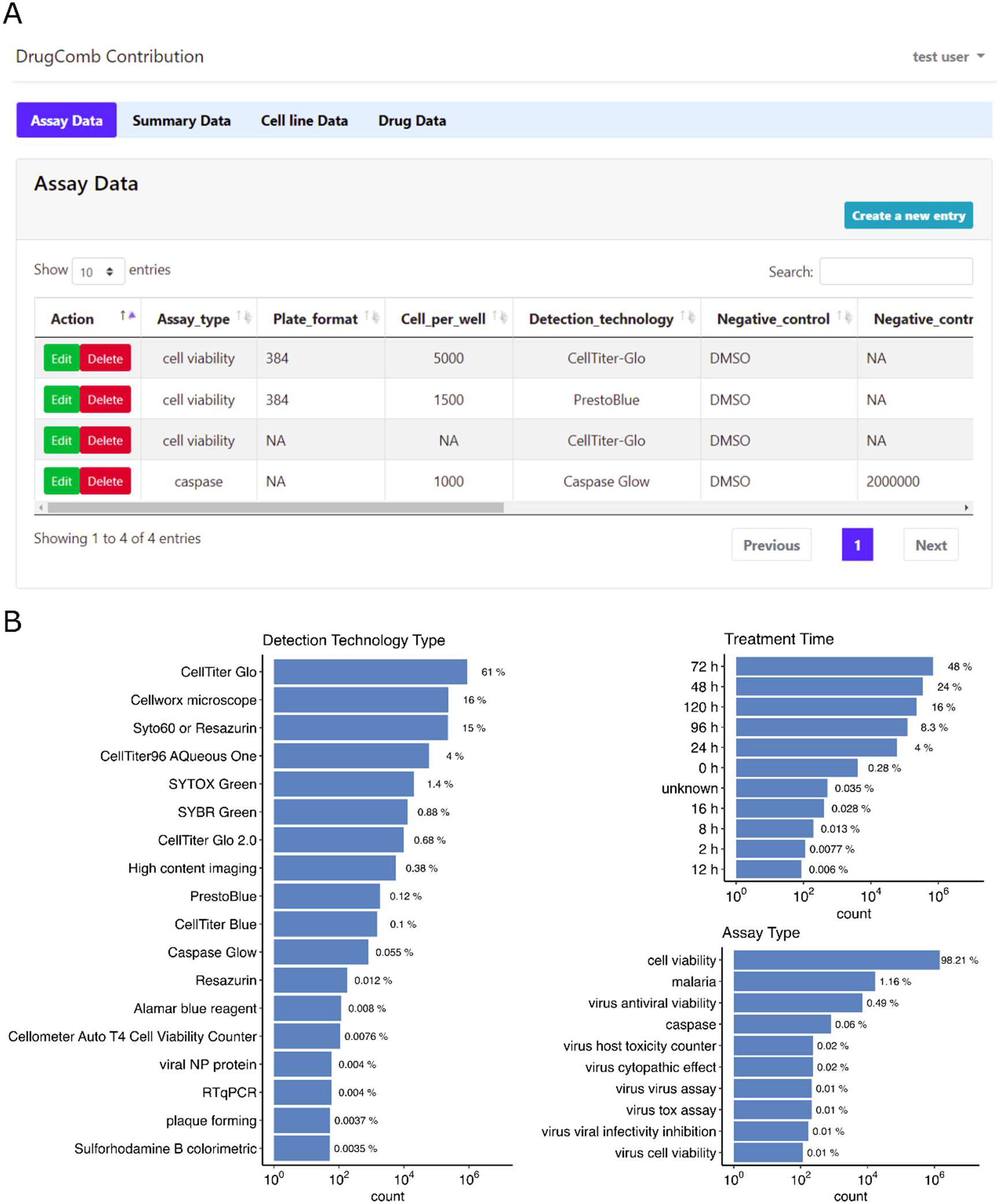
(A) The data contribution interface enables a community-driven data curation effort. (B) Statistics about assay protocols.

### Technical aspects

DrugComb is built using PHP 7.4.14 [Laravel Framework 6.20.7] for server-side data processing, Javascript ECMAScript 2015 for the frontend, D3.js 5.7.0, Vis.js 4.18.1 and Plotly library 1.40.0 for the generation of the interactive visualizations. Data is stored in MariaDB 10.3.17 with RMariaDB 1.0.6.9000 as the driver for interfacing with R. Software development tools including Python 3.6.7, numpy 1.14.1, pandas 0.23.4, scikit-learn 0.20.2, RDkit 2018.03.4, R version 3.5.1, synergyfinder 1.8.0 and tidyverse 1.2.1 are used in the analytical pipelines. Linux distribution CentOS-8 with the kernel 4.18.0 64-bit running on four processor cores and 64 Gb of RAM is used for hosting the web service on a computational cluster.

The data portal has been designed in a straightforward manner to maximize the user flexibility to retrieve the existing datasets as well as to analyse their own datasets. We provided the API access at http://api.drugcomb.org such that users can request data as json files. The API is implemented using the PHP laravel framework. Instructions of each of the modules are provided in their associated web pages and the overview of the data portal was summarized as tutorial video available at the home page. We aim to continue accommodating new features such as cloud-based computing and data infrastructure to facilitate the FAIRness (Findable, Accessible, Interoperable and Reusable) of drug screening data analysis. Meanwhile, the community-based features such as data contribution and quality control can be developed further.

## DISCUSSION

Making cancer treatment more effective is what a combination therapy aims to achieve. With the advances of high-throughput drug screening technologies, an increasing number of drug combinations have been tested. However, before we can develop robust machine learning and network modelling algorithms to predict and understand the potential drug combinations, the datasets need to be systematically curated and harmonized. Here we report the major updates of DrugComb, a comprehensive data portal for the drug discovery community to access the concurrent high-throughput drug combination as well as monotherapy drug screening datasets. These datasets have been deeply curated, standardized and harmonized with the data analysis tools including synergy and sensitivity scoring, such that their potential can be maximized within a unified framework. Furthermore, we have updated the network modelling of the drug combinations, such that the transcriptomics profiles of the cancer cell lines and drug target profiles can be integrated in a signalling network where the protein-protein interactions may provide deeper insights on the mechanisms of drugs and drug combinations. In addition, we have provided a machine learning model to predict a given drug combination for a cell line at the single dose level. To the best of our knowledge, this is the first drug combination prediction tool that has been made online with easy accessibility for drug discovery users.

The four basic modules of DrugComb, i.e. 1) data curation 2) synergy and sensitivity scoring, 3) network modelling, and 4) machine learning constitute a work flow of network pharmacological approaches based on which we may gain deeper understanding of drug-drug interactions. Further steps of DrugComb will involve constant improvement on the data coverage, for example, by including the recent technique of microfluidic-based drug screening (45), as well as the use of more patient-derived samples such as 3D organoid-based drug screening and patient-derived xenograft mouse models. The informatics tools for analysing drug combinations will be made openly accessible such that a consensus on the most appropriate methods can be achieved. We also envisage the high-quality data in DrugComb shall serve as a golden resource for the development of robust and predictive machine learning models, as well as accurate network-based models to capture the mechanisms of drug combinations that may eventually lead to predictive biomarkers that warrant patient stratification for maximizing the efficacy of combinatorial therapies.

## AVAILABILITY

The synergy and sensitivity scores in DrugComb are freely available for download. Larger batch download of raw data are permitted upon agreement on the data sharing policy. The AstraZeneca drug combination datasets are proprietary and a separate agreement is needed. The visualization results for sensitivity, synergy and network models are downloadable as images. The source code for analysing the drug combination datasets is available as the R package SynergyFinder version 2.2.4 (https://bioconductor.org/packages/release/bioc/html/synergyfinder.html). We are committed to open data and welcome any researchers to participate in the development of data curation and harmonization tools for drug discovery.

## SUPPLEMENTARY DATA

Supplementary Data are available at NAR online.

## ACKNOWLEDGEMENT

We thank the authors of the drug combination studies to share their datasets, especially the AstraZeneca Open Innovation program for providing the Dream Challenge data to be part of DrugComb. We thank also the DepMap consortium and the Cell Model Passports to make the transcriptomics profiles of cancer cell lines freely available. We thank the NCATS and other institutions for making their drug screening datasets easily accessible. The data portal is located at the CSC- IT Center for Science in Finland.

## FUNDING

This work was supported by the European Research Council (ERC) starting grant DrugComb (Informatics approaches for the rational selection of personalized cancer drug combinations) [No. 716063]; European Commission H2020 EOSC-life (Providing an open collaborative space for digital biology in Europe [No. 824087]; Academy of Finland Research Fellow grant [No. 317680]; and the Sigrid Jusélius Foundation. Funding for open access charge: ERC starting grant DrugComb.

## CONFLICT OF INTEREST

None declared.

## Notes

https://drugcomb.org/

### Competing Interest Statement

The authors have declared no competing interest.

